# Metamorphism of catalytic domain controls transposition in Tn3 family transposases

**DOI:** 10.1101/2022.02.23.481423

**Authors:** Alexander V. Shkumatov, Nicolas Aryanpour, Cédric A. Oger, Gérôme Goossens, Bernard F. Hallet, Rouslan G. Efremov

## Abstract

Transposons account for a remarkable diversity of mobile genetic elements that play the role of genome architects in all domains of life. Tn3 is a family of widespread and among first identified bacterial transposons notorious for their contribution to dissemination of antibiotic resistance. Transposition within this family is mediated by a large TnpA transposase facilitating both transposition and target immunity. The structural framework for understanding the mechanism of the TnpA transposition is however absent. Here, we describe the cryo-EM structures of TnpA from Tn4430 in apo form and paired with transposon ends. We show that TnpA has a unique architecture and exhibits a family-specific regulation mechanism involving metamorphic refolding of the RNase H-like catalytic domain. The TnpA structure, constrained by a double dimerization interface, creates a unique topology that suggests a specific role for target DNA in transpososome assembly, activation, and target immunity.

## Introduction

Through their ability to move and rearrange DNA sequences, transposons represent an inexhaustible source of genetic innovations^1,2^, such as *de novo* creation of genes, the establishment of regulatory networks, or the exchange of genetic material by horizontal transfer^3,4^ and, in bacteria, emergence and spread of novel antibiotic resistances^5–8^. Among these, Tn3-family transposons were the earliest bacterial transposons to be identified owing to their implication in the transmission of ampicillin resistance^9^. Since then, members of the Tn3-family were isolated from virtually all bacterial groups where they act as mobile platforms for a variety of passenger genes including those conferring resistance to multiple classes of antibiotics^8,10^ (**Supplementary Fig. 1**). Notably, Tn3 family transposons are involved in the recent outbreak of carbapenem-resistant enterobacteria and dispersal of resistance to colistin, which compromises the use of these two “last-resort” antibiotics^11–15^.

Central to the efficiency of these transposons is their replicative “paste-and-copy” transposition mechanism, which duplicates the transposon along with its integration into the target DNA^10,16^. Transposition is initiated by the transposase TnpA, an unusually large member (~1000 amino acids) of the DDE/D superfamily of nucleotidyl transferases. TnpA cleaves and joins the 3’-ends of the transposon to the target using a conserved RNase H-like domain^10,17–19^ (**Supplementary Fig. 2**). This generates a strand transfer product that is then processed by the host replication system, producing two copies of the transposon (**Fig. 1a**). The reaction proceeds through the formation of a unique nucleoprotein complex, transpososome, which brings together the whole donor molecule carrying the transposon and the target (**Fig. 1a**). This distinguishes the “paste-and-copy” mode of transposition from the “cut-and-paste” and “copy-out-paste-in” mechanisms used by other DDE/D transposases during which the element is fully detached from the donor prior to its integration into the target^16,20,21^. TnpA is the only known transposon-encoded protein that facilitates both transposition and target immunity^10^, an intriguing regulation mechanism that prevents multiple transposon insertions into the same target and is thought to prevent self-destruction^22–28^.

**Fig. 1.**
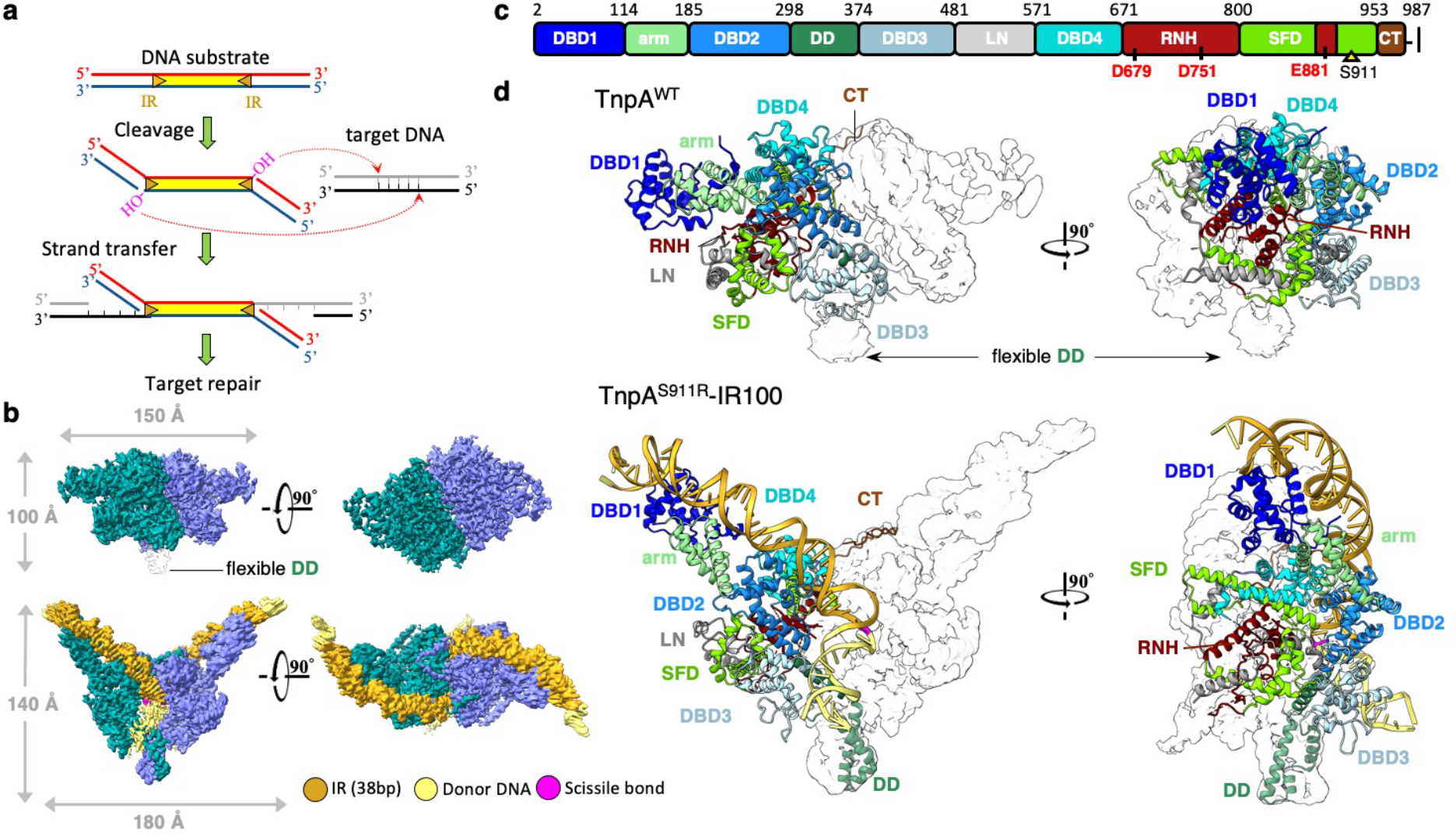
Architecture of TnpA in apo and PEC conformations. **a**, Schematic drawing of “paste and copy” transposition mechanism. **b,** Cryo-EM reconstructions of TnpA^WT^ apo and TnpA^S911R^-IR100 PEC complexes. **c,** Linear diagram of structural domains constituting TnpA. Positions of the conserved catalytic triad DDE and activating mutation are indicated. **d,** Fold and subdivision of TnpA into structural domains is shown in a cartoon representation for a monomer in apo and PEC conformations shown in panel **b**. Domains are color coded as in panel **c**.

Despite their importance, the transposition mechanism of Tn3-family transposons remains poorly understood. Recent studies on the Tn3-family member Tn4430 have started to unravel this mechanism thanks to the characterization of gain-of-function TnpA mutants defective in target immunity^28,29^. However, in the absence of structural information, the molecular interpretation of the data remained very sketchy. Here, we determined single-particle cryo-EM structures of Tn4430 TnpA from *Bacillus thuringiensis* in the apo state and in complexes with transposon ends mimicking the TnpA-transposon complex before and after DNA strand transfer.

## Results

### Structure determination

Cryo-EM structure of wild type TnpA, TnpA^WT^, expressed in *Escherichia coli* was solved to an average resolution of 3.6 Å (**Fig. 1, Table 1, Extended Data Figs. 1,2**). Our attempts to obtain a paired end complex (PEC)^29^ comprising TnpA^WT^ and two Tn4430 terminal inverted repeats (IR) resulted in only a minor fraction of TnpA-DNA complexes (**Extended Data Fig. 2b**). Therefore, we used previously identified hyperactive and immunity deficient mutant TnpA^S911R 28,29^; this enabled us to solve the structures of PEC with two linear IR substrates IR100 and IR48, and one branched substrate IR71st, prefiguring the strand transfer product of an IR end into the target DNA. The substrates contained 38 base pair (bp) long TnpA recognition sequences of the IR end ^10^ (**Extended Data Fig. 1a**). The structures were solved to a resolution between 2.9 and 3.1 Å (**Extended Data Figs. 3-5, Table 1**). The poor density of the flexible N-terminal domain was improved by multibody refinement^30^ (**Extended Data Fig. 4**), allowing *ab initio* modeling of 92% of the 987 residue-long protein and up to 54 bp long DNA substrate (**Supplementary Table 1**).

**Table 1.**
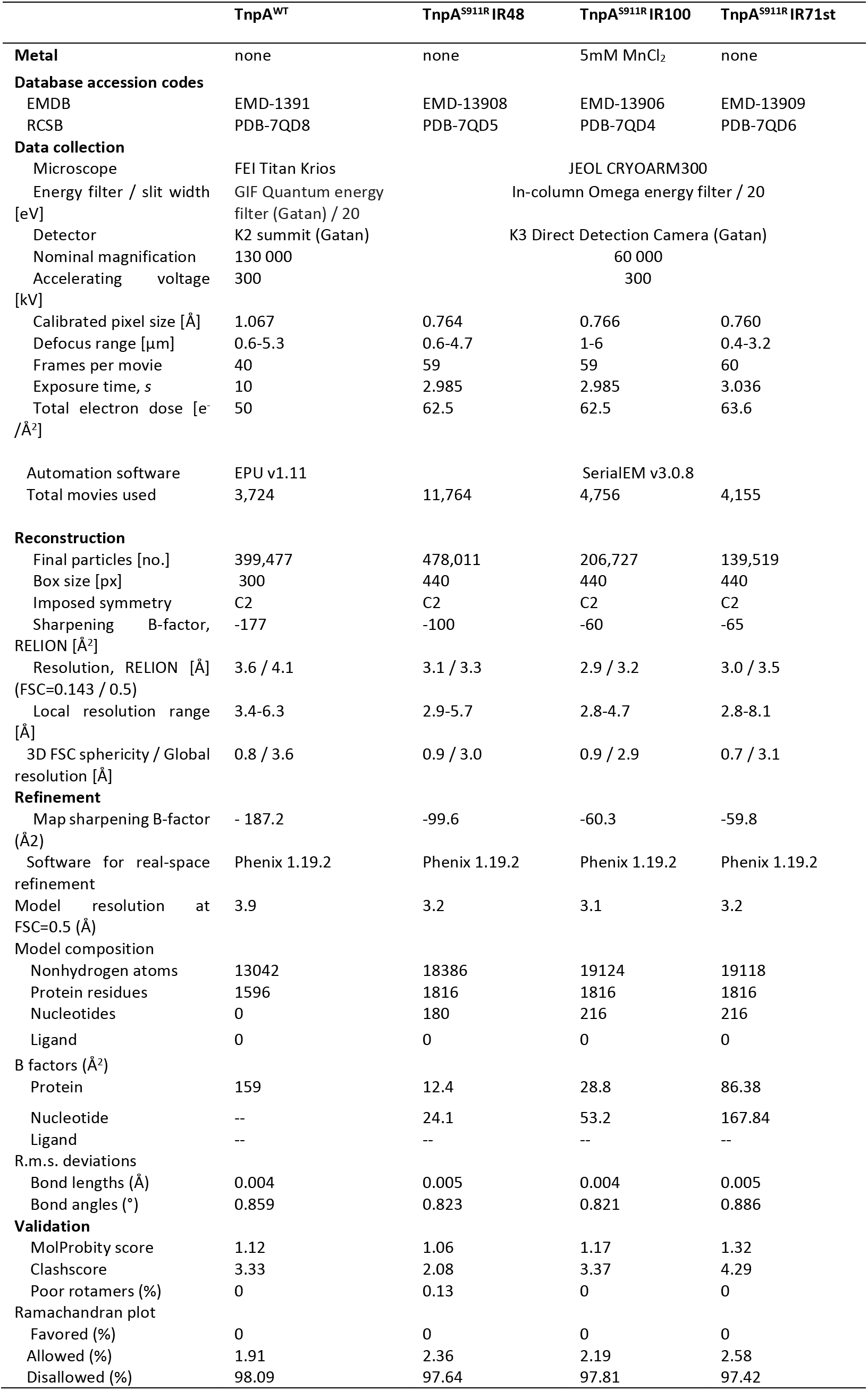
Statistics of cryo-EM data collection and model refinement.

### Architecture of TnpA

In both apo and PEC conformations, contrary to a previously proposed model^29^ TnpA exists as a dimer (**Fig. 1b**). Binding of transposon ends is accompanied by large conformational changes that transform the compact apo form into an expanded V-shaped PEC with approximately 140 Å long edges (**Fig. 1b,c, Supplementary Video 1**). The bound IR sequence curves smoothly while the outer flanking sequence (OFS) bends sharply at the level of TnpA cleavage site (**Figs. 1,2a**). The scissile phosphate is positioned at the center of the dimer, where it faces the RNase H-like domain of another TnpA protomer. This cross-reactivity mechanism, wherein one subunit recognizes a specific DNA sequence (cis-interaction) while another catalyzes end cleavage and strand transfer (trans-interaction) is a convergent feature of most characterized DDE/D transposases despite their structural heterogeneity^21,31–34^.

**Fig. 2.**
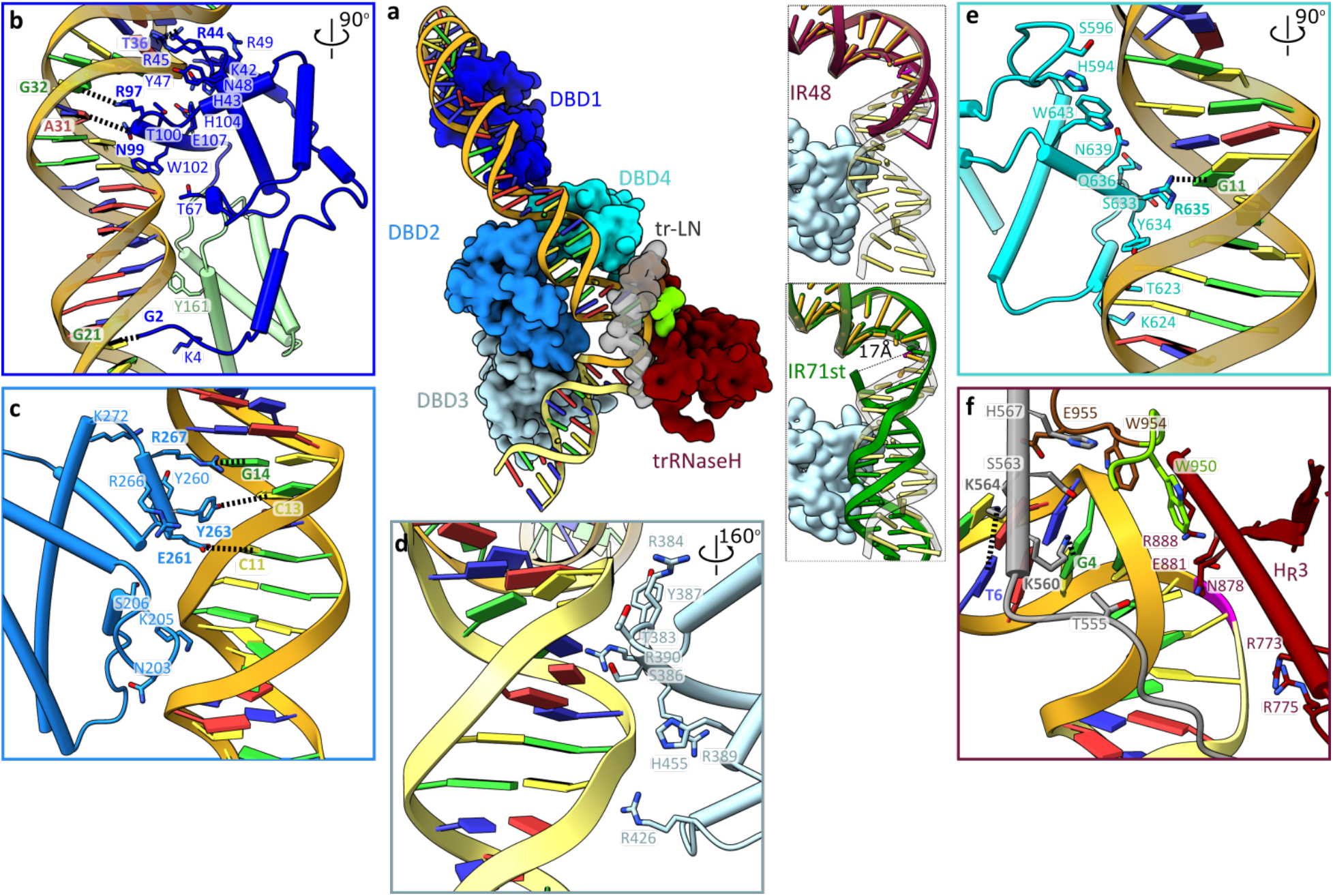
TnpA-DNA interactions. **a,** Domains interacting with DNA substrate IR100. Insets display conformations of OFS for IR48 (purple, top) and IR71st (green, bottom). **b-f,** Details of DNA interaction with DBD1(**b)**, DBD2 (**c)**, DBD3 (**d)**, DBD4 (**e)**, and trans interactions (**f)**. Base-specific interactions are shown by dotted lines and corresponding residues are labeled in bold.

Each TnpA protomer can be divided into ten structural domains (**Fig. 1c,d**), most of which are α-helical. They are arranged in a 140 Å long stem and a discoidal-shape protrusion with 50 Å diameter in the center. The stem is composed of four DNA-binding domains (DBD), an α-helical arm domain that separates DBD1 from DBD2, DBD3, and DBD4 by approximately 40 Å, and a dimerization domain (DD) positioned at the end of the stem (**Figs. 1d, 2a**). The protrusion is made of a long linker (LN) bridging DBD3 and DBD4 and a catalytic RNase H-like (RNH) domain encircled by α-helices which we refer to as a scaffold domain (SFD). SFD is composed of a structurally unique RNase H insertion domain^21,31,32,34–36^ and a protein sequence downstream of RNH (**Fig. 1c, Supplementary Video 1**). Overall, the architecture of TnpA is unique, essentially constituted of novel folds.

Protein-protein dimerization interfaces are very similar between apo and PEC conformations and are associated with two distinct areas within each protomer: DD and a C-terminal tail (CT). DD protrudes out of the dimer (**Fig. 1d, Supplementary Video 1**) as an extended α-helical bundle, in which three tightly packed α-helices contributed by each protomer form primarily hydrophobic interactions (**Extended Data Fig. 6a**). In the apo form, DD is flexible, but low-resolution density suggests that the interaction between DD is unchanged (**Fig. 1d**).

The 30 residues long CTs interlock the protomers by docking conserved residues onto the surface of the adjacent protomer (**Extended Data Fig. 6b**). In the PEC, the dimer is further stabilized through interactions with DNA, which also stabilizes CT-mediated dimerization (**Fig3a**, **Extended Data Fig. 6b**). This unusual dimerization interface that locks the TnpA dimer at both extremities has important consequences for the transposition mechanism.

**Fig. 3.**
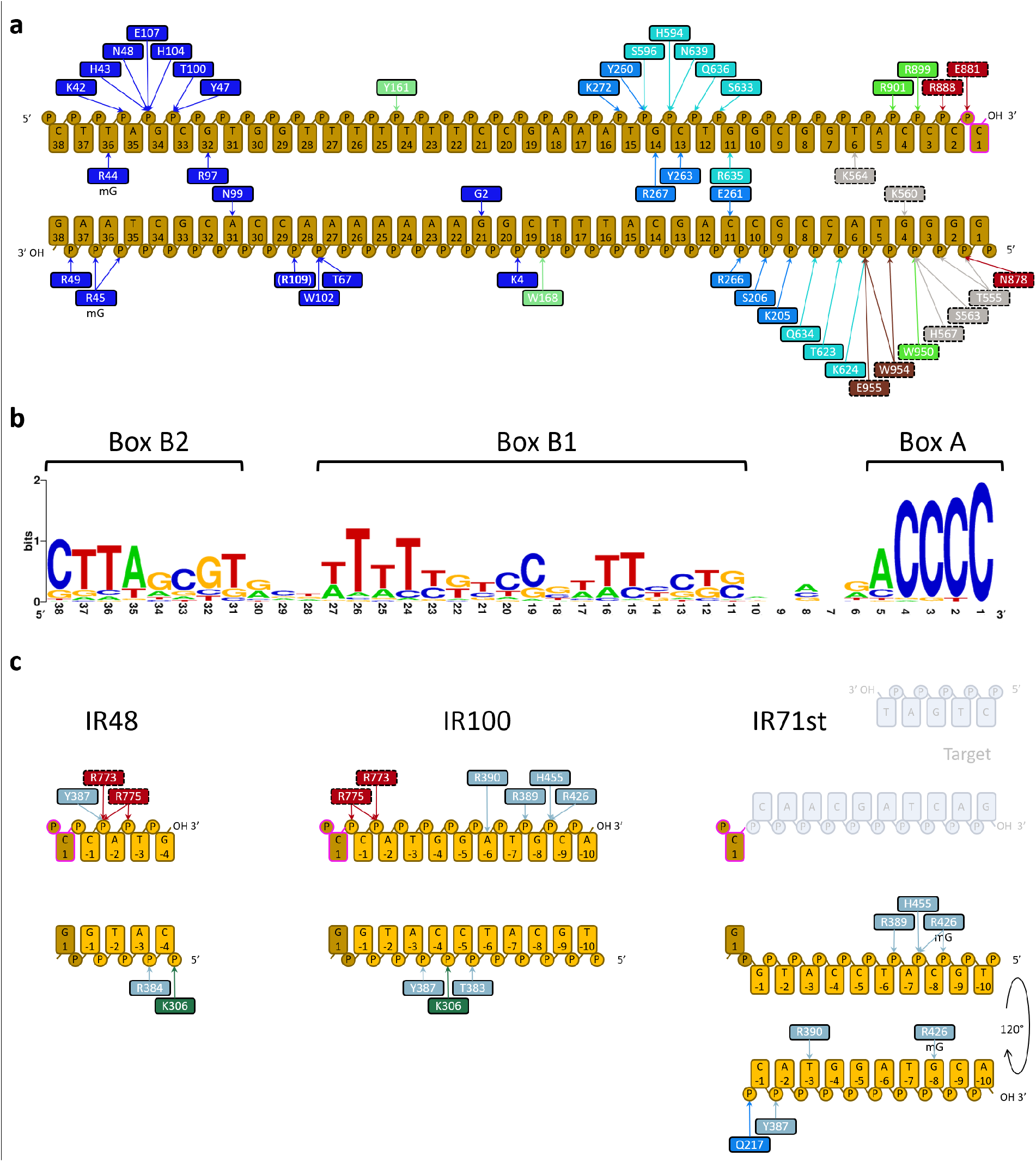
Schematic representation of TnpA-DNA interactions and conservation of the recognition sequence. **a,** Interaction of TnpA with recognition sequence. **b**, Conservation of the recognition sequence. Three conserved regions named Box A, Box B1 and Box B2 are indicated. **c**, DBD3-OFS interactions for IR48, IR100, and IR71st constructs. In panels **a** and **c** residues are color coded by domain color as in **Fig. 1d**. Trans interactions are shown with dashed borderline around the corresponding box.

### Interaction with transposon ends

An extended 130 Å long positively charged DNA-binding surface (**Extended Data Fig. 6c,d**) creates over 50 polar contacts with a 47 bp long cis-DNA (**Figs. 2,3**) that includes the 38 bp IR sequence and a 9 bp OFS from the donor locus (**Figs. 1,2**). The structural differences between the three TnpA-DNA complexes are confined to the differences in the conformation of the OFS fragment (**Figs. 2a,3c, Supplementary Video 2**).

DNA remains base-paired throughout its length, but OFS bends sharply by approximately 72 ° at the trans-DNA interaction site next to the DNA cleavage site (**Figs. 1d, 2a,f**). The bending site corresponds to the highly conserved box1 of the recognition sequence (**Fig. 3b**) and is likely primarily caused by electrostatic interactions between the OFS and DBD3 (**Fig. 2d**). This suggestion is supported by the reduction in the bending angle to 54 ° for the IR48 substrate, in which the interaction of OFS (5 bps) with DBD3 is reduced (**Fig. 2a,3c, Supplementary Video 2**). This DNA binding and bending pattern is consistent with DNA footprint analysis showing extended protection of the OFS together with hypersensitive sites around the TnpA cleavage sites in PEC^29^.

DNA bending creates stress that is released upon DNA cleavage. Thus, in the post strand transfer IR71st substrate with a broken transferred strand (**Extended Data Fig. 1a**), the OFS is rotated by approximately 120 °around G0, separating ^C1^C5 from ^C-1^C3 by 17 Å (**Figs. 2a,3c, Supplementary Video 2**). Similar to the bending of the target DNA^32,35,36^, bending of the OFS likely helps in making the transposition irreversible.

DBD3 binds OFS non-specifically. DBD3-OFS interaction is restricted to the interaction with with DNA backbone (**Fig. 2d**) at positions −1 to −8. The interaction is also promiscuous as, permitting DNA binding to DBD3 in different positions and orientations (**Figs. 2a,3c, Supplementary Video 2**). Here again, the establishment of a different set of contacts between TnpA and the OFS before and after strand transfer may stabilize the strand transfer product and favorizing the transposition reaction in the forward direction.

DBD1, 2, and 4 interact with the recognition sequence in a sequence-specific manner (**Figs. 2b,c,e, 3a**). DBD1 is one of the key determinants of specificity. It unexpectedly shares fold similarity and DNA-binding surface with BEN domains, a new class of DBDs found in a variety of transcriptional factors involved in chromatin silencing and gene repression in eukaryotes^37^ (**Extended Data Fig. 7a**). DBD1 interacts with conserved DNA sequence of box B2 and forms more than 20 polar interactions spread over 50 Å long contact surface (**Figs. 2a,b, 3a,b**). It interacts specifically with bps in both the minor and major groove between positions 21 and 36 (**Fig. 2b**).

One DNA helical turn down from the cleavage site (bps 6-15), DBD2 and DBD4 are docked into a major groove of IR (**Fig. 2c,e**). DNA sequence recognition is mediated by a short α-helix (residues 261-267) on DBD2 (**Fig. 2c**) and residue R635 on DBD4 (**Fig. 2e**) and occurs with nucleotides C11, G11, C13 and G14 at the beginning of box B1 that have a low level of conservation (**Fig. 3a**). DBD4 is structurally homologous and shares a DNA-binding mode with the N-terminal domain of Tn5 transposase^31^ (RMSD 2.7 Å over 53 residues), despite only 9% sequence identity (**Extended Data Fig. 7b**).

TnpA trans-DNA interactions are mediated by the linker, SFD, and RNH domains with bps between positions 1 and 6 that includes the highly conserved box A, viz. 5’GGGGT (**Figs. 2f, 3a,b**). Base-specific interactions with LN domain K560-G4 and K564-T6 contribute to DNA sequence recognition.

Consistent with the high specificity of TnpAs for their respective IR^10,28^, sequence recognizing residues and corresponding nucleotides, with exception of R44-T36 pair, display modest or no conservation (**Supplementary Figure 2**). Therefore, conservation of the transposon recognition sequence^10^ likely reflects the geometric constraints required for matching the DNA backbone to the extended DNA binding surface of TnpA, while the sharp DNA bend between recognition sequence and OFS occurs at the highly-conserved sequence of box A, at the very end of the transposon. Here sequence conservation likely reflects the requirement for and mechanistic importance of OFS bending^38^.

### Conformational changes

Upon apo to PEC transition, the protein module upstream of DBD4 translates and rotates by approximately 50 ° as a rigid body, resulting in a shift of DBD1 by a distance of 40 Å (**Figs. 1, 4a, Supplementary Video 3**). Conformational changes are a prerequisite for the tight binding of IR and assembly of PEC. It renders the surface of DBD3, otherwise occluded by the linker domain, accessible for DNA binding (**Supplementary Video 3**), and it rearranges DBD2 relative to DBD4 to form the DNA-binding site (**Fig. 4b**). The conformational transition also creates a 30 Å opening, which is absent in the apo conformation, between the bodies of protomers and DD (**Extended Data Fig. 6c,d, Supplementary Videos 1,3**).

**Fig. 4.**
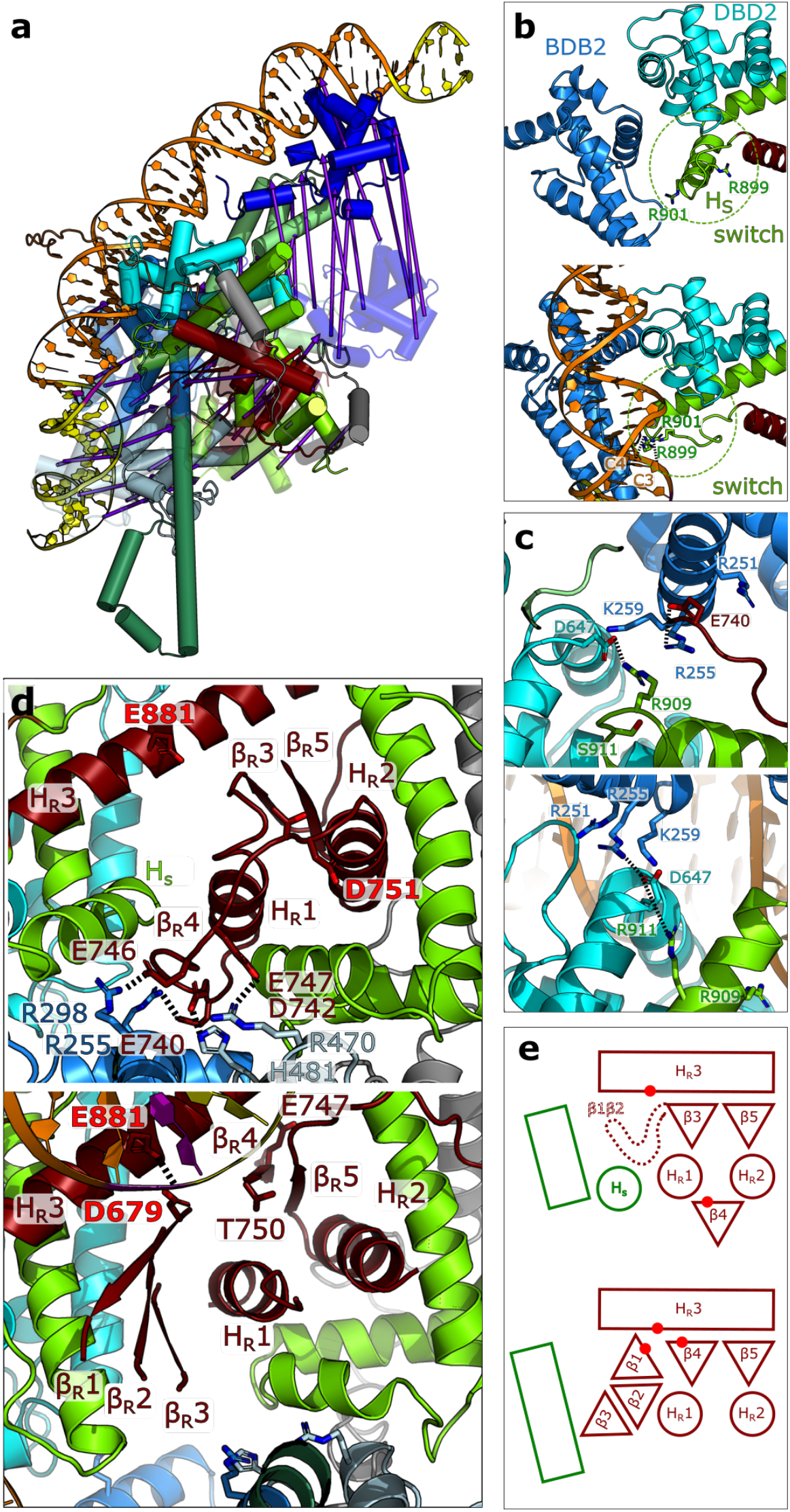
Conformational changes between apo and PEC states. **a,** Structures of apo (semitransparent) and PEC conformations aligned to invariant C-terminal region. Arrows indicate movements of the domains from apo to PEC states. **b-e,** Details of conformational changes. Top and bottom panels correspond to apo and PEC state, respectively. **b**, Rearrangement of DBD2 relative to DBD4 and refolding of switch helix H_s_. **c**, Ionic lock, and salt bridge R911-D647 stabilizing the PEC state. **d,e**, Metamorphic behavior of RNH domain fold. In **d** the catalytic residues are labeled in red. In **e** rectangles and circles correspond to α-helices, triangles to β-strands. Red dots indicate the positions of the catalytic residues.

The apo to PEC transition involves a peculiar conformational change in the conserved region of the SFD. A helix (residues 897-908), further referred to as the switch helix, H_S_, changes its fold. The N-terminal turn of H_S_ (residues 897-901) unfolds in a loop, while the rest refolds into an extension of a long scaffold helix. This local refolding reorients R899 and R901, buried in the apo state, towards the protein surface where it interacts with the cis-DNA backbone in positions C4 and C3, respectively (**Figs. 3a,4b, Supplementary Video 3**).

Upon transition to PEC several salt bridges found in a cavity proximal to S911 and connecting different domains are disrupted (**Fig. 4c**). The S911R mutation introduces an additional positive charge into the cavity, which likely destabilizes the electrostatic interactions and facilitates apo to PEC transition. This hyperactive mutation further stabilizes PEC by forming an R911-D647 salt bridge (**Fig. 4c**). Mapping other hyperactive and target immunity deficient mutations^28^ on the TnpA structure showed that most of them destabilize the apo conformation (**Extended Data Fig. 8**), suggesting that the apo-PEC transition controls both the activity and target immunity of the transposase.

### Metamorphic RNase H-like domain

The conserved fold of the RNase H superfamily, three α-helices (H_R_1-3) flanking a five-stranded ß-sheet (ß 1-5)^39^, is consistent with the TnpA secondary structure prediction^28^. However, the RNH is among the least ordered parts of the complex and displays an unusual metamorphic behavior (**Fig. 4d-e**). In the apo conformation, adjacent α-helices H_R_1 and H_R_2 sandwich a short 2-stranded β-sheet (β3, β5) with catalytic H_R_3, while predicted β4 is positioned on the opposite side of the α-helical pair, where it is stabilized by salt bridges with DBD2 and DBD3 (**Fig. 4c, Extended Data Fig. 9a**). The helical scaffold tightly wrapped around the RNH and specifically H_s_ precludes assembly of the 5-stranded β-sheet (**Fig. 4d**), and consequently, the density for β1 and β2 is not observed in the apo conformation. In such a non-canonical conformation, RNH is partially disassembled while the DDE catalytic site is completely disorganized (**Fig. 4d,e**).

In PEC, refolding of H_s_, together with rearrangement of the scaffold, allows for the folding of the RNH β-sheet which now appears as a low-resolution density consistent with an extended β-sheet which was modelled with the aid of AlphaFold2^40^ (**Fig 4d, Extended Data Fig. 9b,c**).

Among the catalytic triad residues, only E881 on H_R_3 is rigidly positioned and faces C1 (**Figs. 2f, 4d**), whereas D679 and D751, despite being predicted to be closely positioned, are very mobile, independent of the presence of divalent ions (**Table 1**). This is consistent with the low *in vitro* TnpA activity^29^ and suggests that additional factors are required to activate the catalysis.

Protein metamorphism has been described as a regulatory strategy in several proteins^41^ but to the best of our knowledge, it has not been observed for RNase H domains and thus presents a unique regulatory mechanism among the Tn3 family transposases.

TnpA displays structural similarity to the core of the cut-and-paste Tn5 transposase^31^ (**Extended Data Fig. 7c**). However, in Tn5, the SFD is missing, and consequently neither switch helix nor metamorphic refolding is observed^31,42^. This indicates a common origin but a remarkably different evolution of TnpA and Tn5 transposases, which are mechanistically distinct.

## Discussion

Models for paste-and-copy replicative transposition mediated by Tn3-family transposons and other bacterial elements were among the first ones to be proposed in the literature^43^ and are presented as a classical mechanism of transposition in textbooks. This mechanism has been described with great molecular details for bacteriophage Mu which uses replicative transposition to multiply its genome during lytic development^44^. However, the relevance of the Mu paradigm for non-viral elements such as Tn3-family transposons is questionable. Mu transposition is mediated by two main proteins: the transposase MuA and the target binding protein MuB also involved in transposase activation and target immunity. For the Tn3-family, these different functions are fulfilled by a single, non-related transposon-encoded protein, the transposase TnpA. Hence, as shown here for Tn4430, the molecular architecture of Tn3-family transpososome, and the mechanisms that control its assembly and activity are unique and totally differ from those of Mu^35^.

Unlike cut-and-paste transposition that excise the transposon from the donor molecule by the formation of double-strand breaks at both ends, or the copy-out-paste-in mechanism during which replication generates a circular copy of the transposon that can diffuse away prior to integration into a new locus^16,20,21^, initiation of paste-and-copy transposition is a one-step process. It requires the assembly of an elaborate transposition complex in which two distant regions of the genome the donor and the target, are brought together to catalyze single-strand DNA cleavage and joining reactions between the transposon ends and the target DNA (**Fig. 1a**) ^10,16,44^ These reactions must be highly concerted and regulated because incomplete or abortive transposition can damage both the donor and the target molecule, thus compromising the survival of the host cell and therefore that of the transposon. In return, being coupled to DNA replication, paste-and-copy transposition is likely one of the most powerful mechanisms to promote dispersal of passenger genes and to bring about specific DNA rearrangements such as deletions, inversions and replicon fusions that have been shown to play a crucial role in bacterial genome evolution, notably by reassorting multidrug-resistant plasmids in response to antibiotherapy pressure^45,46^.

Cryo-EM structures reported here reveals that active Tn4430 transpososome assembly is indeed controlled at multiple levels and point toward a crucial role of target DNA in the activation mechanism and its plausible link with target immunity.

Consistent with previous biochemical data^29^, the structural signatures of the TnpA^S911R^ PEC suggest that it represents a transpososome-like complex without the target DNA. The distance of ~30 Å between the symmetry-related scissile bonds is consistent with 5-bp staggered insertion of the transposon ends into the target DNA^10^. The opening between the dimerization domains is sufficiently large to accommodate double-stranded DNA, its surface is positively charged (**Extended Data Fig. 6d**) and highly conserved when compared to the rest of the TnpA surface (**Extended Data Fig. 6e**). Despite being mobile, the RNase H-like domain is assembled and correctly positioned to cut the transposon DNA. Furthermore, low-resolution density features of the TnpA^S911R^-IR71st reconstruction revealed position of the target-mimicking branch of the substrate next to the trans-RNH domain (**Fig. 5a**). This indicates that the surface of RNH domain exposed to the cavity has affinity for DNA. The target DNA bound within the active site may in turn stabilize the RNH in its active conformation. The angle of approximately 90 ° between the target-mimicking DNA branches is consistent with the target DNA bending commonly observed in transposases^32,35,36^.

**Fig. 5.**
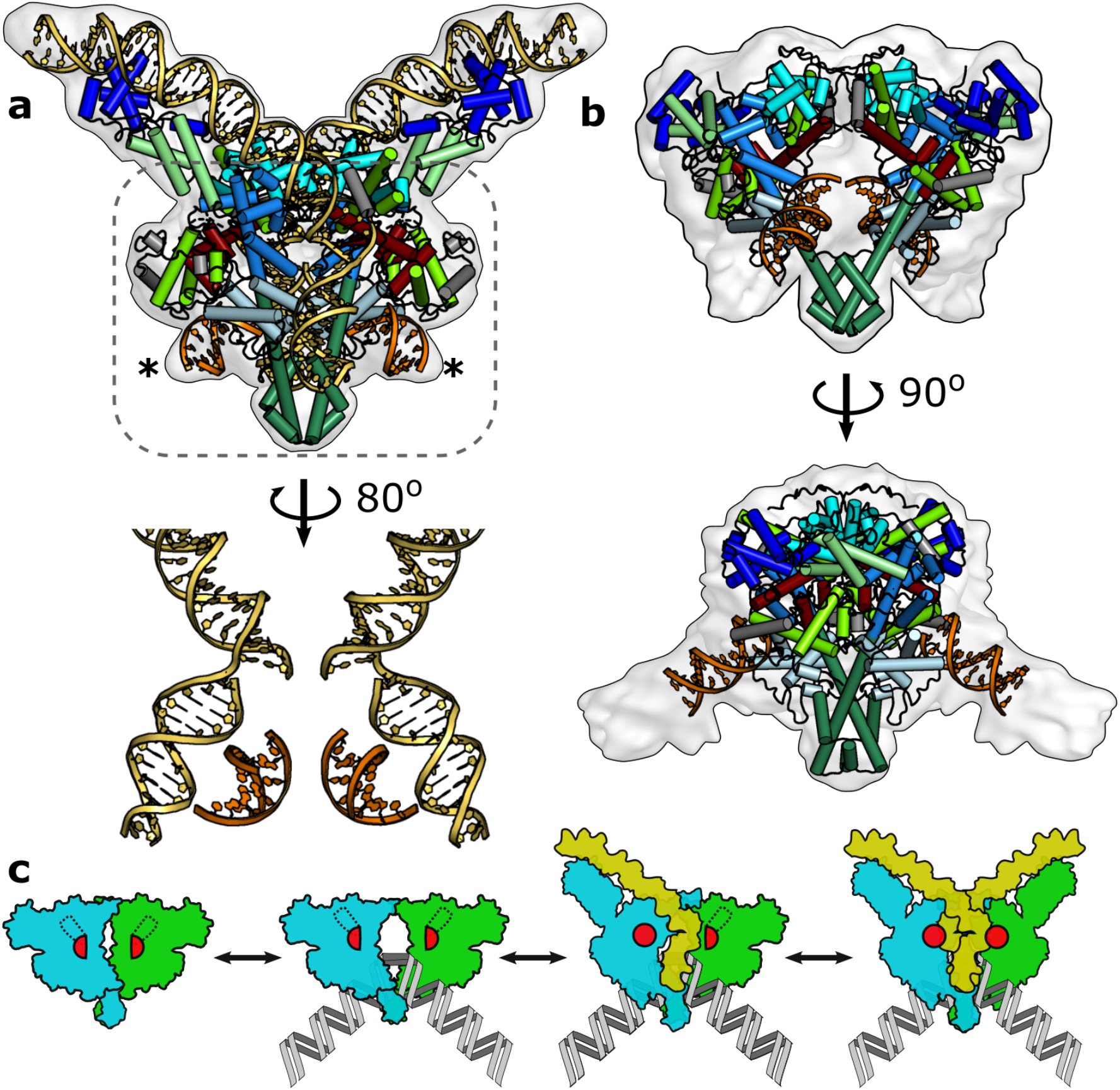
Plausible target DNA binding sites and non-activated DNA-bound apo state point towards mechanism of transpososome assembly. **a,** Reconstruction of TnpA^S911R^-IR71st reveals additional low-resolution density consistent with the target DNA branch bound to trans SFD and RNH domains (orange, marked with an asterisk). **b,** TnpA^WT^-IR100 conformation with bound DNA shown in orange displays the opening between the dimerization domains. **c,** Cartoon of the proposed mechanism for transpososome assembly. The RNase H domain is shown by a red (semi) circle, and the target DNA is shown in grey.

The double dimerization interface that closes the TnpA dimer is stabilized in PEC. This suggests that the target DNA either threads through the opening observed in PEC conformation as a double-stranded break or more likely, that it binds prior to the formation of PEC. The dataset of TnpA^WT^ was collected in the presence of IR100 substrate. Even though a majority of particles are found in DNA-free state, a smaller fraction of particles revealed TnpA^WT^-IR100 complex for which low-resolution reconstruction was obtained (**Fig. 5b, Extended Data Fig. 2b,c**). Unexpectedly, the structure of TnpA^WT^-IR100 rather than correspond to single-end complex (SEC) in which one transposon end is bound specifically to TnpA^29^, revealed a conformation intermediate between apo and PEC in which DBD1 and DBD2 are in an apo-like conformation while the rest of the protein is in a PEC-like conformation. Ends of the straight DNA fragments are bound to DBD3, mimicking the position of OFS in PEC. This conformation of TnpA^WT^-IR100 complex indicates that the TnpA dimer is flexible and does not need to be fully activated to create the opening for target DNA binding between the protomers (**Fig. 5b**). The SEC conformation was not unambiguously resolved although 2D class averages with features expected for SEC were observed in TnpA^WT^-IR100 dataset (**Extended Data Fig. 2b, red box**) indicating that SEC might be present in the ensemble albeit at low occupancy.

Taken together, our structural data suggest a plausible Tn3-family specific transposon assembly and activation mechanism, schematically depicted in **Fig. 5c**. First, the target DNA binds. The binding might be enabled by transient disruption of dimerization interface mediated by CT fragments and occur spontaneously or with assistance of other unknown factors that specifically direct transpososome assembly to permissive target. Binding of target DNA is followed by the sequential binding of transposon ends. Upon binding, cis switch helix refolds and triggers folding of the cis RNH concomitant with transition first to SEC and consequently to active PEC conformation.

Tethering of TnpA on target DNA creates an avidity effect which may contribute to the mechanism of target immunity. Tethered TnpA would sample adjacent sequences of the DNA with higher probability than those located on separate DNA molecules or on remote regions of the target DNA. Thus, whenever transposon is already present in the target, TnpA would preferably bind to transposon recognition motif on the same DNA molecule and outcompete other transposons by forming an unproductive complex. Alternatively, binding of TnpA to transposon ends might promote the dissociation of TnpA-target complexes in cis, in a same DNA locus, thereby forcing active transpososome to only assemble on a remote site. Whichever the mechanism, the requirement for TnpA unlocking as revealed by the structures reinforces the idea that transposition can only be effective when all partners including the two transposon ends and an appropriate target assemble in the complex.

## Conclusions

Here, we have described the structures of the Tn3-family transposase in apo and PEC conformations. The structures of TnpA reveal a common cross-activation mechanism but unusual metamorphic behavior of the switch helix and the RNase H-like domain, pointing toward a novel regulatory mechanism among DDE/D transposases. The double dimerization interface presents an unprecedented feature of TnpA and suggests a well-defined sequence of events involving both the donor and target DNA to assemble of the active transpososome. The resolving structure of a completely assembled TnpA transpososome will be the next important step toward understanding the transposition mechanism with atomic details.

## Materials and Methods

### DNA substrates

IR substrates were generated by annealing specific oligonucleotides (**Supplementary Table 2, Extended Data Fig. 1a,b**) at 95°C for 10 min, followed by cooling to room temperature.

### Protein production, purification, and characterization

Tn4430 TnpA^WT^ and TnpA^S911R^ mutant were fused to a cMyc-His6 epitope tag at the C-terminus and were expressed in *E. coli* TOP10 cells under the control of the pAra promoter^29^. Cells were grown at 37°C in TB media containing tetracycline (12.5 ug/ml) till OD600 reached 0.7-0.8; the temperature was then dropped to 18°C. To induce the cellular chaperones, benzyl alcohol (0.1%) was added, and the cells were grown for 2 h before induction with L-arabinose (0.04%). After 3-4 h the medium was topped with L-arabinose (0.12%) and cells were grown overnight. The following day, the bacteria were centrifuged at 4°C (7,000 *g*, 45 min), the pellet from 500 ml of bacterial culture (~ 5 g) was re-suspended in 20-30 ml of buffer A [50 mM HEPES (pH 7.9), 1 M NaCl, 10% glycerol, and 20 mM imidazole] supplemented with cOmplete EDTA-free inhibitor cocktail tablet (Sigma-Aldrich), and flash frozen in liquid nitrogen. The thawed cell suspension was supplemented with 0.25 mg/ml lysozyme (Sigma-Aldrich), 0.1% Triton-X (Sigma-Aldrich), MgCl_2_ (10mM), and DNase I (Sigma-Aldrich). The mixture was diluted to a final volume of 20 ml using buffer A and incubated for 1 h at 4°C on a rotator. After sonication, the lysate was cleared by centrifugation (18 000 *g*, 45 min), filtered through a 0.45 um filter, and supplemented with 5 mM ATP and 4 mM MgCl_2_ before loading on a 5 ml HisTrap column (Amersham) pre-equilibrated in buffer A. Next, the bond material was washed with two column volumes (CV) of buffer A containing 0.1% Triton-X and two CV of buffer A containing 5 mM ATP and MgCl_2_ interspersed by washes with buffer A. TnpA was eluted with a 20–500 mM linear gradient of imidazole in buffer A over 16 column volumes. Pooled fractions were concentrated by ultrafiltration (Amicon Ultra-15, 100 K MWCO), and applied to a Superose 6 Increase 10/300 GL column equilibrated in 50 mM HEPES (pH 7.9), 200 mM NaCl, and 100 mM L-Arg HCl. The fractions containing pure TnpA were pooled and mixed with 4- to 10-fold molar excess of DNA substrates. The mix was kept at 4°C on a rotator either for several hours (TnpA^WT^) prior to vitrification or overnight (TnpA^S911R^) followed by concentration (Amicon Ultra-15, 100 K MWCO) and gel filtration to remove unbound DNA using Superose 6 10/300 GL (GE Healthcare) column equilibrated in 50 mM HEPES (pH 7.5), 100 mM NaCl, and 30 mM L-Arg HCl. Freshly purified TnpA-DNA complexes were used directly for the preparation of cryo-EM grids.

The homogeneity and oligomeric state of the apo and complex forms were assessed using mass photometry on a Refeyn OneMP instrument (Refeyn Ltd.), which was calibrated using an unstained native protein ladder (NativeMark™ Unstained Protein Standard A, Thermo Fisher Scientific Inc.). Measurements were performed at concentrations of 0.1-0.2 mg/ml using AcquireMP 2.2.0 software and were analyzed using the DiscoverMP 2.2.0 package (**Extended Data Fig. 1d**).

### Preparation of cryo-EM grids

The grids for the TnpA^WT^-IR100 complex were prepared by mixing freshly purified TnpA at a concentration of 0.5 μM with a 4-fold excess of IR100 substate, incubated for 1 h on ice, and applied to a Quantifoil holey carbon grid (R2/1, 300 mesh) glow-discharged in an ELMO glow discharge system (Cordouan) at 0.3-0.35 mBar and current of 10-15 mA for 60 s. Grids were blotted from one side for 3 s at 70-90 % relative humidity and plunge-frozen in liquid ethane using a Cryoplunge 3 System (Gatan).

TnpA^S911R^ samples with DNA substrates were applied to graphene-oxide-coated grids. The aqueous dispersion of graphene oxide (GOgraphene; William Blythe Ltd) was diluted in doubledistilled water (ddH2O) to a final concentration of 1.3 mg/ml, followed by gentle sonication in Elmasonic S 30 (H) for 120 s in a cold room and spun down at 300 g for ~2 min. C-flat holey carbon grids (R2/1, 300 mesh) were glow-discharged as described above, and 4 μl of GO solution was applied on the grids, followed by one minute incubation; subsequently, the GO solution was removed by blotting briefly with Whatman No.1 filter paper and washed by applying 20 μl ddH2O onto the graphene-oxide-coated side twice and once on the back side of the grid with blotting steps in between. TnpA^S911R^-DNA substrate complex (5 μl) at a final concentration of 0.13-0.16 mg/ml was applied on a GO-covered grid, blotted from one side for 3 s at ~90 % relative humidity, and plunge-frozen in liquid ethane using a Cryoplunge 3 System (Gatan).

### EM data acquisition

The TnpA^WT^ was imaged at the CM01 beamline at ESRF^47^ using the EPU v1.11 software for automated data acquisition on a Titan Krios cryo-electron microscope (Thermo Fisher Scientific) operated at 300 kV equipped with a Quantum LS electron energy filter (Gatan). Image stacks were recorded with a K2 Summit (Gatan) direct electron detector operating in counting mode at a recording rate of 4 raw frames per second. The microscope magnification was 130,000 (corresponding to a calibrated sampling of 1.067 Å per pixel). The total dose was 50 electrons per Å^2^ with a total exposure time of 10 s, yielding 40 frames per stack. A total of 3724 image stacks were collected with a defocus range of 0.6 - 5.3μm (see **Table 1** for details).

Micrographs for TnpA^S911R^ complexes with DNA were collected at 300 kV on a CRYO ARM 300 (JEOL) electron microscope at a nominal magnification of 60,000 and corresponding pixel size of approximately 0.76 Å. The images were recorded using a K3 detector (Gatan) operating in correlative-double sampling (CDS) mode. The microscope illumination conditions were set to spot size 6, alpha 1, and the diameters for the condenser and objective apertures were 100 and 150 μm, respectively. The energy filter slit was centered on the zero-loss peak with a slit width set to 20 eV. Coma-corrected data acquisition^48^ was used to acquire between 6 and 25 micrographs per stage position using SerialEM v3.0.8^49^. Each micrograph was recorded as a movie of 59 or 60 frames over a 3 s exposure time and at a dose rate of 11 e^-^pixel^-1^s^-1^ (corresponding to a dose rate per frame of 0.6 e^-^Å^-2^) and total exposure dose of approximately 60 e^-^Å^-2^ (see **Table 1** for details).

### Image processing

Initial data processing was performed on-the-fly using RELION_IT^50^. Dose-fractionated movies were subjected to motion correction and dose weighting using the MotionCorr2^51^. The dose-weighted aligned images were used for CTF estimation using CTFFIND-4^52^. An in-house script was used to plot the calculated parameters, visualize the results, and select the micrographs for further processing (Shkumatov et al; in preparation). The aligned and dose-weighted images were imported into cryoSPARC v3.1.0 ^53^, and CTF was calculated using Patch CTF. Particle selection was performed using a blob or template-based picker followed by several rounds of 2D classification. An *ab initio* reconstruction and initial 3D refinement were performed using cryoSPARC. The 2D classification of the TnpA^WT^ dataset in cryoSPARC revealed four different populations of classes, including higher-order oligomers (**Extended Data Fig. 2b**, blue frame). The different conformations were further separated by *ab initio* model calculations and heterogeneous refinement. Separated subsets were independently reconstructed by applying homogeneous and non-uniform refinement. (**Extended Data Fig. 2**). For the processing of TnpA^S911R^ datasets, particles were imported into RELION 3.1^50^. The low-pass filtered to 60 Å initial model was used for 3D auto-refinement using C1 symmetry. This was followed by multiple rounds of 3D-refinement and 3D classification using either C1 or C2 symmetry, CTF refinement, and Bayesian polishing^54^ (**Extended Data Figs. 3-5**). To improve the density corresponding to the N-terminal domain in TnpA^S911R^-DNA complexes, the signal for the monomer was subtracted, followed by multibody refinement using two rigid bodies^50^ (**Extended Data Fig. 4**). Local resolution was estimated in RELION 3.1, with a B-factor from the postprocessing job. The directional resolution of the final map was measured using the 3DFSC server^55^.

### Model building and refinement

Initially, parts of the model were built automatically using the PHENIX v1.19.1 map_to_model procedure^56^. This was followed by a manual model building in COOT 0.9.5^57^. Poorly resolved β-strand of the RNase H domain in TnpA^S911R^-DNA complexes were predicted using AlphaFold2^40^ and fitted into the density as a rigid body with a scaffold domain. Next, regions for which density was absent were removed from the model. The models were refined with the PHEINIX v1.19.2 real_space_refine procedure^58^ against maps filtered using the local filter procedure of RELION 3.1. Secondary structure, Ramachandran, and ADP restraints were applied during the refinement procedure that included ‘global_minimization’ and ‘local_grid_search’ strategies. For the TnpA^S911R^-IR71st complex during the last iteration, ADP restrains were relaxed. The models were validated using MolProbity^59^. The model and data statistics are reported in the **Table 1**. The TnpA^WT^-IR100 model was constructed by first fitting refined TnpA^S911R^-IR100 in the low-resolution density map followed by real space refinement of the model in COOT with applied ProSMART restraints using the initial TnpA^S911R^-IR100 complex as a reference model. The resulting structure was not refined further because of the low resolution of the map.

### Visualization and Sequence alignment

The images of proteins were prepared using PyMol v2.4.2 and ChimeraX v1.2.4^60^ programs. Sequence alignments were performed in Clustal Omega^61^ and visualized with ESPript 3.0 server^62^. The phylogenetic tree was generated with the help of the TnCentral database^63^.

## Acknowledgments

We are indebted to Dr. Adam Schröfel and Dr. Marcus Fislage for their assistance with cryo-EM data collection, and to Dr.Gipsi Lima-Mendez for her precious assistance and expertise in phylogenetic and bioinformatic analyses. We acknowledge the European Synchrotron Radiation Facility for providing the beam time on CM01, and we would like to thank M. Hons for assistance. This work benefited from access to the Netherlands Centre for Electron Nanoscopy (NeCEN) at Leiden University with the assistance of Dr. C. Diebolder and Dr. J. Ortiz Espinoza. We thank Prof. S. Raunser & Dr. O. Hofnagel for providing access to the electron microscope at the Max Planck Institute for Molecular Physiology. We thank the VIB Tech Watch fund for facilitating access to the Refeyn instrument. We would like to acknowledge the funding provided by Vlaams Instituut voor Biotechnologie, Fonds Wetenschappelijk Onderzoek (Grant Nos. G0H5916N, G054617N to R.G.E), and Fonds Spéciaux de Recherche (UCLouvain) and Fonds National de la Recherche Scientifique (CDR grants J.0162.16 and J.0096.20 to B.F.H).

## Author Contributions

A.V.S., C.A.O, G.G., N.A., and B.F.H. designed the strategy and DNA substrates for the preparation of cryo-EM samples. A.V.S. prepared cryo-EM grids, collected EM data, and processed cryo-EM data. R.G.E built and refined the models. All authors have analyzed the results. R.G.E prepared the original draft of the manuscript. A.V.S., B.F.H., and R.G.E. reviewed and edited the manuscript. A.V.S, N.A and R.G.E prepared the figures. All authors have analyzed and discussed the structures. All authors have discussed and edited the manuscript. B.H. and R.G.E. conceived, managed, and supervised the project and acquired the funding.

## Data availability

Cryo-EM density maps and atomic models were deposited in the PDB and EMDB databases under the following accession codes: TnpA^WT^ (PDB ID: 7QD8, EMD-13910), TnpA^S911R^-IR100 (PDB ID: 7QD4, EMD-13906), TnpA^S911R^-IR48 (PDB ID: 7QD5, EMD-13908), and TnpA^S911R^-IR71st (PDB ID: 7QD6, EMD-13909).

---

**Extended Data Fig. 1.**
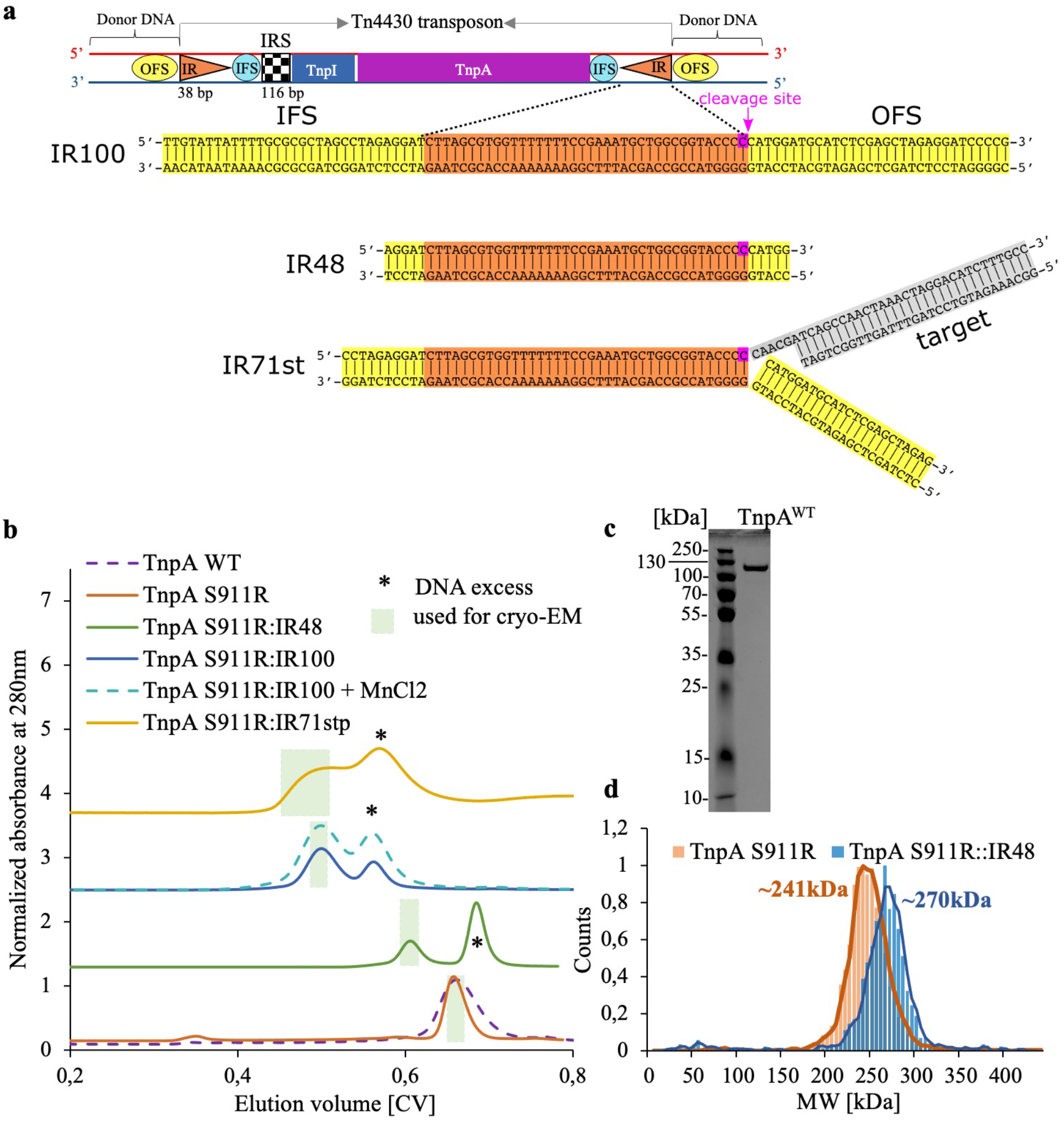
DNA substrates, purification, and characterization of TnpA. **a**, Structure of transposon and DNA substrates used for structure determination IFS: inner flanking sequence; OFS: outer flanking sequence. **b**, SEC profiles of TnpA with and without DNA substrates. The green rectangles correspond to the fractions used for cryo-EM. **c**, SDS-PAGE of purified TnpA. **d**, Mass photometry measurements indicate homogeneous TnpA populations in the apo state and in complex with IR100 substrate.

**Extended Data Fig. 2.**
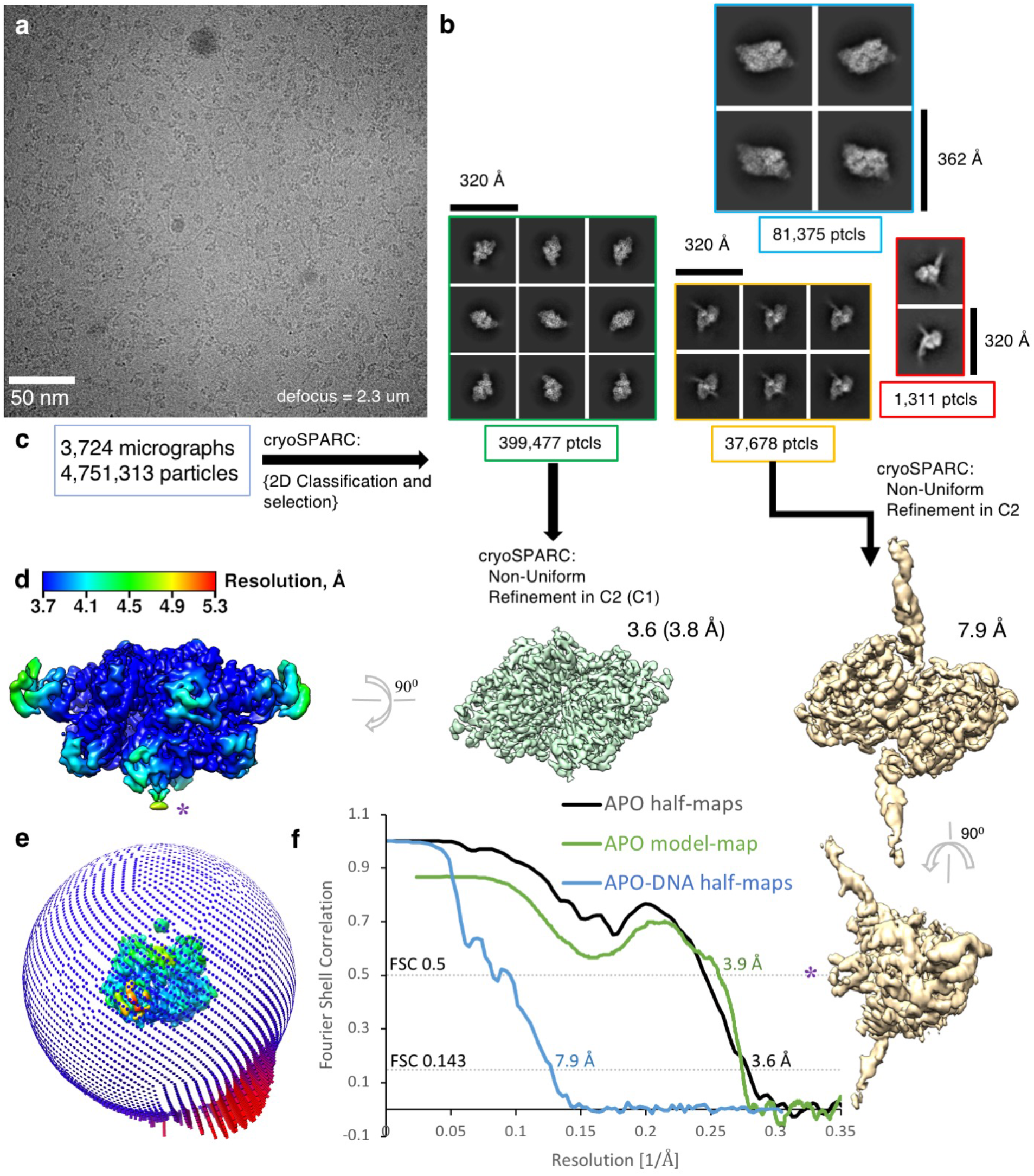
Cryo-EM reconstruction of TnpA^WT^ in the presence of IR100. **a,** Raw cryo-EM image. **b,** 2D class averages revealed several particle populations. **c,** Schematic representation of 3D reconstruction steps. Two reconstructions were determined from the dataset: 3.6 Å reconstruction of TnpA^WT^ apo state and 7.9 Å resolution TnpA^WT^-IR100 complex. **d,** Local resolution of TnpA^WT^ reconstruction. **e,** Distribution of particle orientations. **f,** FSC plots for half-maps, and between the model and cryo-EM map.

**Extended Data Fig. 3.**
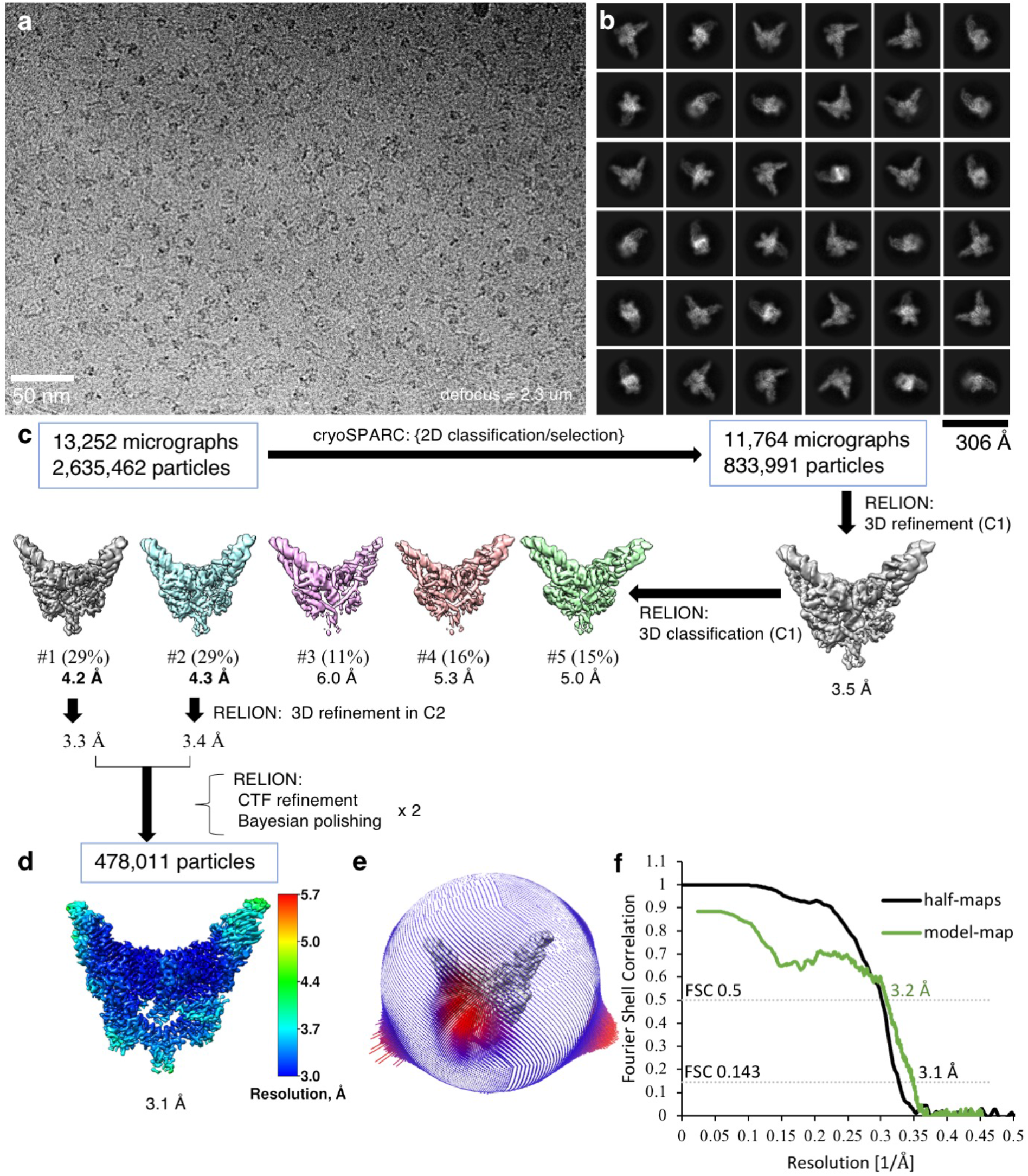
Cryo-EM reconstruction of the TnpA^S911R^-IR48 complex. **a,** Raw cryo-EM image. **b,** 2D class averages. **c,** Schematic representation of 3D reconstruction steps. **d,** Local resolution of the reconstruction. **e,** Distribution of particle orientations. **f,** FSC plots for half-maps, and between the model and cryo-EM map.

**Extended Data Fig. 4.**
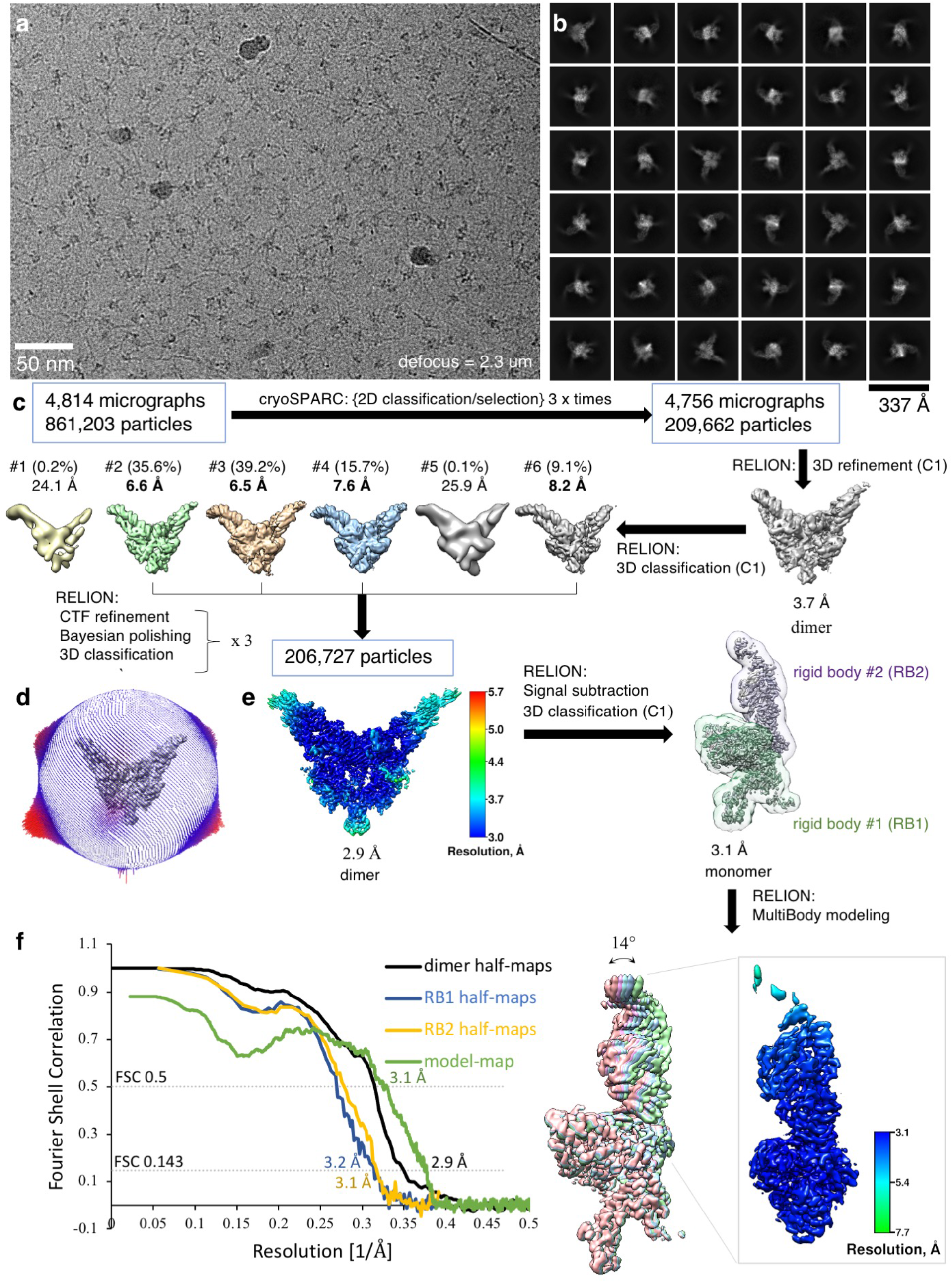
Cryo-EM reconstruction of the TnpA^S911R^-IR100 complex. **a,** Raw cryo-EM image. **b,** 2D class averages. **c,** Schematic representation of the 3D reconstruction steps and properties of the resulting reconstruction. The reconstruction of N-terminal DBD1 was improved by multi-body refinement. **d,** Distribution of particle orientations. **e,** Local resolution of the reconstruction. **f,** FSC plots for half-maps, and between the model and cryo-EM map.

**Extended Data Fig. 5.**
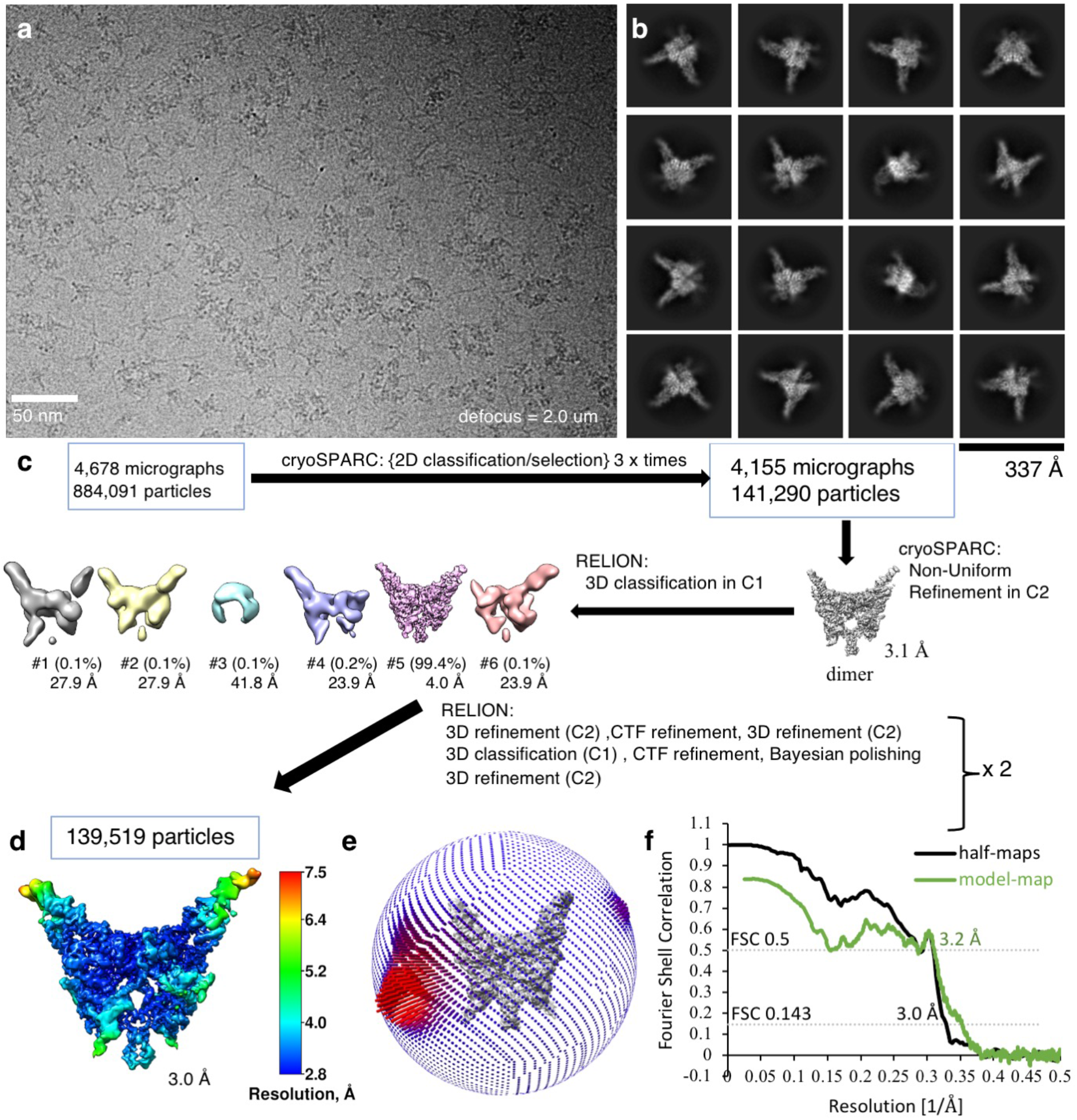
Cryo-EM reconstruction of the TnpA^S911R^-IR71st complex. **a,** Raw cryo-EM image. **b,** 2D class averages. **c,** Schematic representation of 3D reconstruction steps, **d,** Local resolution of the reconstruction. **e,** Distribution of particle orientations. **f,** FSC plots for half-maps, and between the model and cryo-EM map.

**Extended Data Fig. 6.**
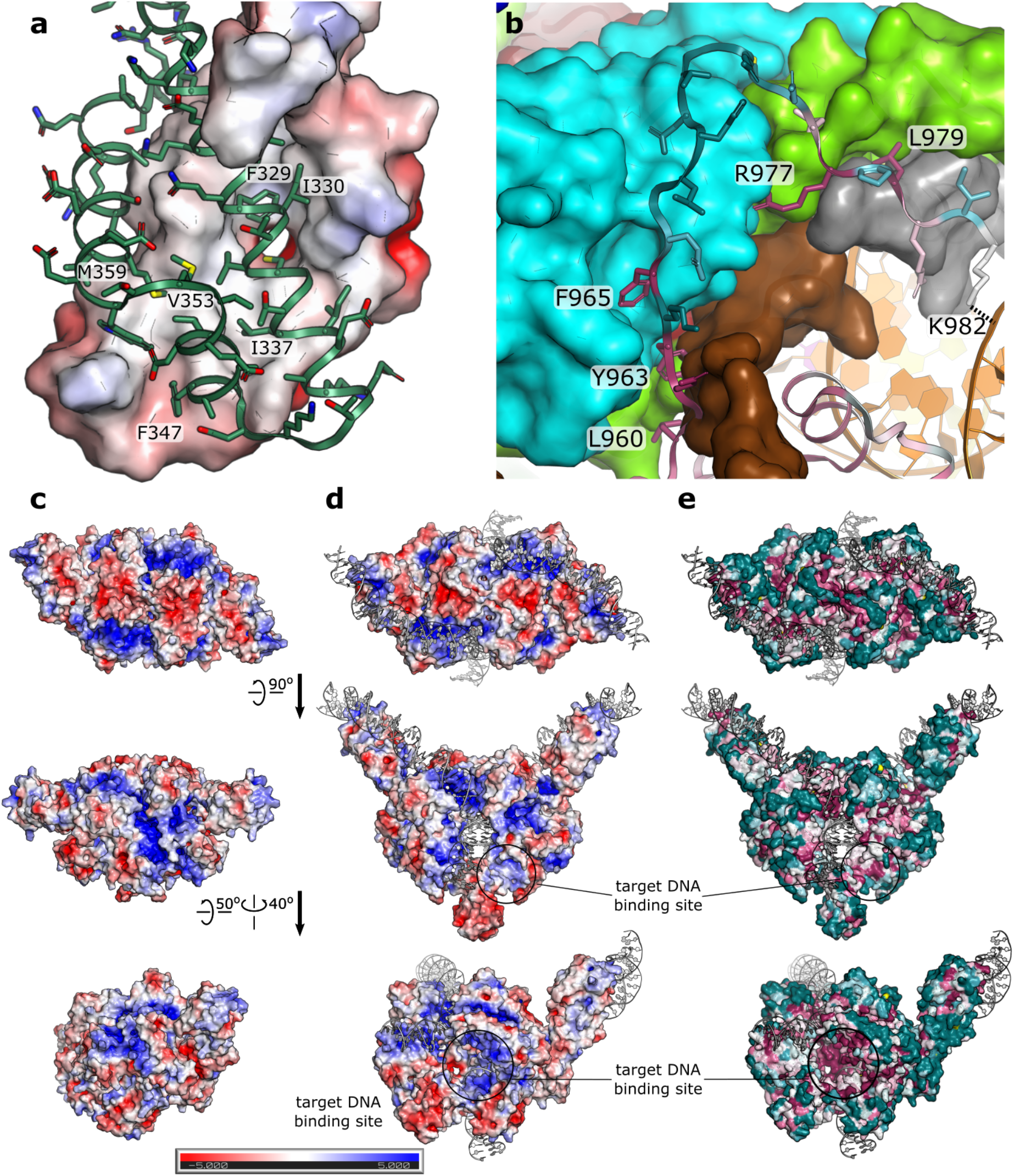
Dimerization interfaces and surface properties of TnpA. **a,** Interaction between DD domains involves nearly exclusively hydrophobic interactions. Surface electrostatic potential is shown for the DD of the second protomer. **b,** Dimerization interactions mediated by the CT. Residues are colored based on the extent of conservation, as calculated by CONSURF^64^. Conservation increases from deep teal to white to purple. The surface of the adjacent protomer is colored by domains color-coded as shown in **Fig. 1d**. **c,** Surface electrostatics for the apo state, and **d,** for PEC. **e**, Conservation of surface residues for PEC conformation as calculated by CONSURF. Protein orientation is same as in panel **d**. Circles indicate surfaces with positive potential and high conservation suggesting putative position for target DNA binding.

**Extended Data Fig. 7.**
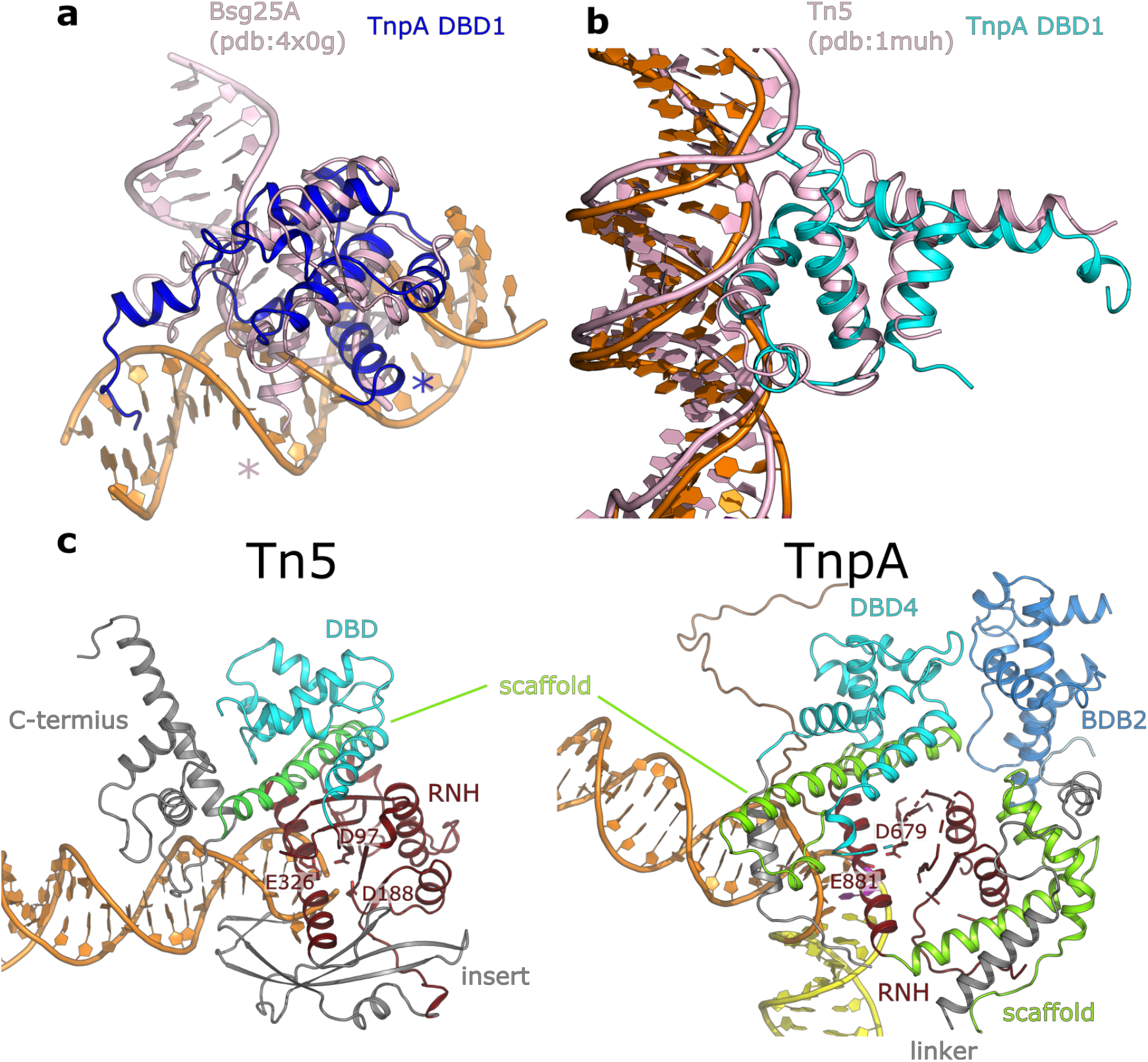
Structural homology of TnpA. **a,** DBD1 displays fold similarity to the BEN domain. Structural overlay is shown with Bsg25A from *Drosophila melanogaster* ^37^; difference in orientation of bound DNA is likely due to difference in the position of DNA-reading helix marked with asterisks. **b,** Structural alignment of Tn5 (PDB identifier 1muh) cis-DNA binding domain to DBD4 of TnpA shows a similar fold and mode of DNA binding. **c**, Similarities and differences in the architecture of Tn5 and TnpA. In Tn5, elements that structurally align with TnpA are color-coded as in **Fig. 1d**, while other domains, including insertion into the RNase H-like domain, are shown in grey. TnpA domains are color-coded as in **Fig. 1d**. Residues of the catalytic DDE tirade are indicated for both proteins. The orientation of the RNase H domains is similar in both proteins. The scaffold that is well pronounced in TnpA is absent in Tn5. For clarity, TnpA domains remote from the homologous core are not shown.

**Extended Data Fig. 8.**
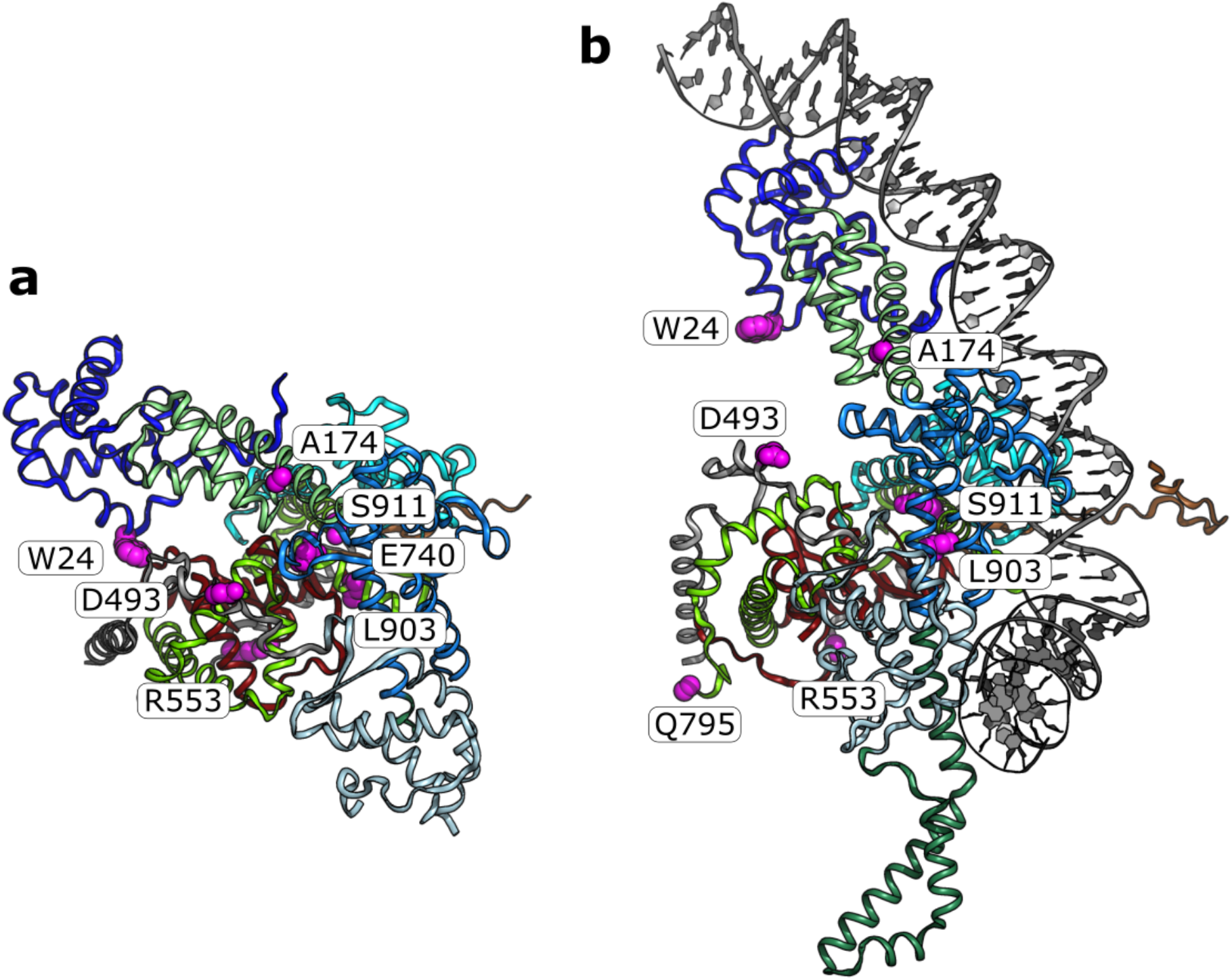
Distribution of hyperactivating mutations. Positions of hyperactive, immunity impairing mutations are shown as pink-sphere representations of the corresponding side chains. **a**, TnpA^WT^ apo conformation and **b,** TnpA^S911R^-IR100 complex. The mutations are located at the interfaces between domains disrupted in the PEC conformation. Only monomers are shown for clarity.

**Extended Data Fig. 9.**
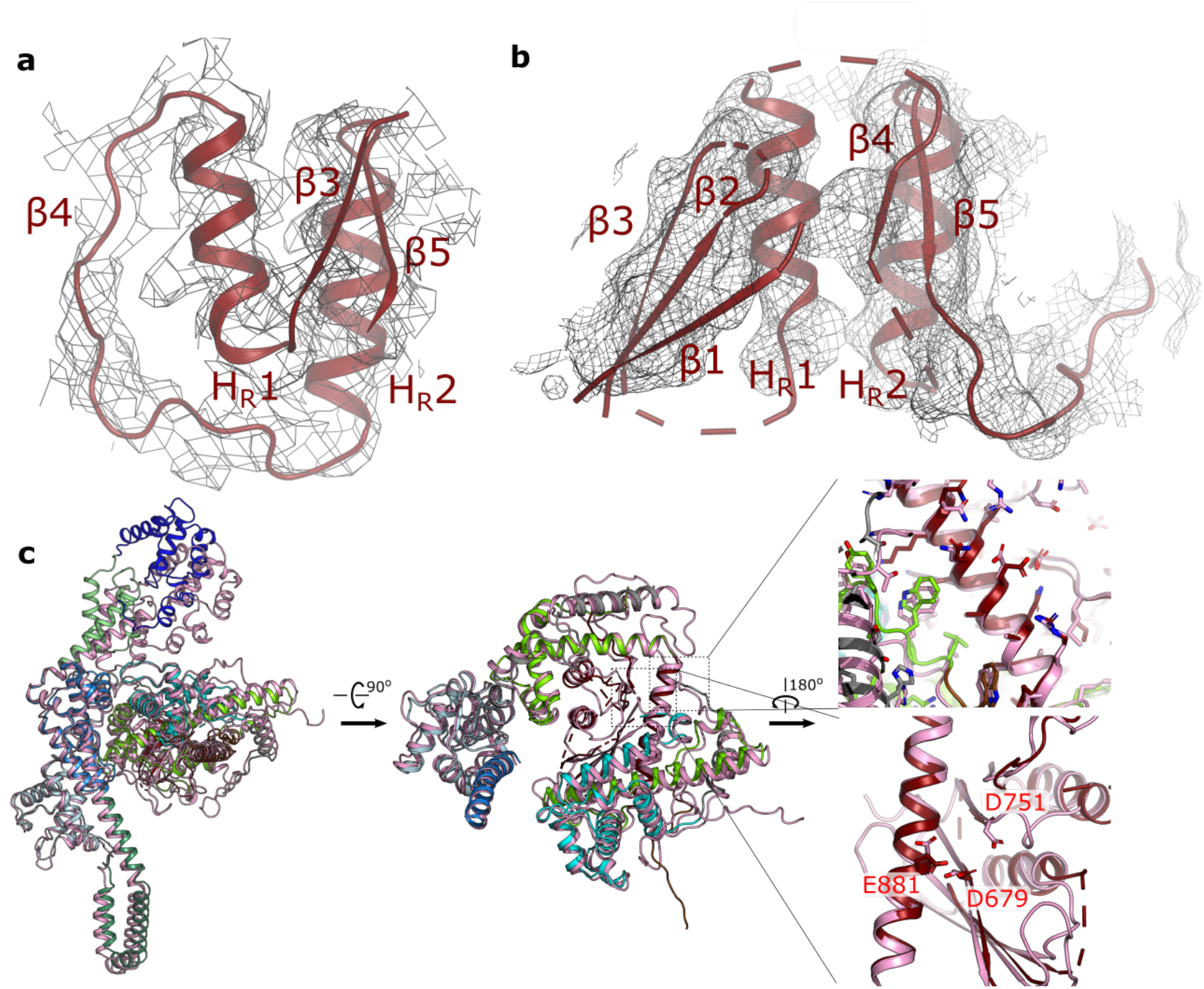
Modelling of the RNH domain and AlphaFold2 model. **a-b,** Models of the RNase H domain in apo (**a**) and PEC (**b**) states and cryo-EM density. **c,** Overlay of TnpA model built into experimental map (without the use of AphaFold2, except bottom right panel) with the AplhaFold2 model. AlphaFold2 prediction of TnpA accurately matched the PEC conformation, albeit deviating at the position of DBD1. The AlphaFold2 model had a high predictive power and accurately traced polypeptides through density regions that were too poor for *ab initio* modeling. This included the 5-stranded β-sheet of the RNase H domain. In the AphaFold2 model the catalytic triad D967, D751, and E881 is appropriately assembled to form an active site.

